## Supplementary information for "Critical Assessment of Metaproteome Investigation (CAMPI): A Multi-Lab Comparison of Established Workflows"

### Contents

1. Supplementary notes 2
1.1 DNA/RNA extraction, processing and sequencing 2
1.2 Database generation 3
1.3. Protein grouping methods 4
1.4. Unipept analyses 6
2. Supplementary Figures 7
2.1 Identification: Peptides and Proteins groups comparison 7
2.2 Protein grouping methods 11
2.3 Taxonomic Analyses: Omics Comparison 15

### 1. Supplementary notes

#### 1.1 DNA/RNA extraction, processing and sequencing

Before sample distribution to all participants, the GUT sample was subjected for an initial DNA extraction and sequencing at the Bielefeld University Center for Biotechnology working group for computational Metagenomics. DNA from the fecal sample was extracted as follows. Briefly, 200 mg of sample was resuspended in 1 ml Tris/HCl (100 mM pH 8.0), supplemented with 100 mM EDTA, 100 mM NaCl, 1% (wt/vol) polyvinylpyrrolidone and 2% (wt/vol) sodium dodecyl sulfate and transferred to a 2 ml Lysing Matrix E tube (Qbiogene, Alexis Biochemicals, Carlsbad, CA). Cells were lysed in a Fastprep-24 instrument (40 s, 6.0 ms-1). Sample was centrifuged at 14.000 rpm for 1 min at 4°C and the supernatant washed with one volume phenol/chloroform (1:1), centrifuged and the aqueous phase washed with one volume chloroform. After centrifugation, nucleic acids (aqueous phase) were precipitated with one volume of ice-cold isopropanol and 1:10 volume of 3 M sodium acetate, incubated during 1h at -20°C and centrifuge during 30 min at 14.000 rpm. Pellet was resuspended in 100 µl of water. The quality and quantity of the DNA samples were analysed on 1% agarose gel and spectrophotometrically by determination of the A260/A280 ratios.

Additionally, to provide meta-omic derived databases for SIHUMIx as well as a deeper sequencing depth for the GUT sample, both SIHUMIx cells pellet and fecal aliquot were subjected to DNA/RNA co-extraction at the University of Luxembourg. There, a dedicated methodological framework that allows for the sequential extraction and purification of all biomolecular fractions from single unique samples was used[1]. Briefly, the snap-frozen fecal samples have been homogenized in a liquid nitrogen bath (6875D Freezer/Mill, SPEX), then aliquoted into 150 mg aliquots for subsequent multi-omics extraction. The extraction is based on the Qiagen Allprep kit (Qiagen) by an automated robotic liquid handling system (Freedom Evo, Tecan). The main advantage of the multi-omics extraction is that DNA, RNA as well as protein extracts are provided by one single aliquot, which facilitates and improves a multi-omics data integration.
For the metagenomic sequencing, 70 ng of DNA was sheared using NGS Bioruptor (diogenode, UCD300) with 30s ON and 30s OFF for 15 cycles. DNA libraries were

prepared using TruSeq Nano DNA kit (Illumina, FC-121-4002) using standard protocol. The libraries were prepared for 350bp average insert size. For the metatranscriptomic sequencing, 500 ng of RNA was rRNA depleted using RiboZero kit (Illumina, MRZB12424). Further library was prepared using TruSeq Stranded mRNA library preparation kit (Illumina, RS-122-2101). Standard protocol was followed for the fragmentation and priming steps as mentioned for the kit.

Prepared libraries were checked using bioanalyzer (Agilient) and quantified using Qubit (Invitrogen). 4 nM pool of the libraries were sequenced on NextSeq500 using 2x150 bp read length.

##### 1.2 Database generation

The bottom-up approach is the standard protocol for LC-MS/MS-based protein identification. By design, it creates a unique problem: the resulting homogenized mixture of proteolytic peptides must be computationally linked back to a specific protein (Hettich et al., 2013). The success of the peptide identification and protein inference is therefore inextricably linked to the quality of the employed database, making its selection and construction one of the most critical steps in a metaproteomic study. Ideally, the database should represent the exact protein and microbial composition of the sample. However, this is not widespread in metaproteomics as sequencing each sample is time- and cost-intensive. Using nonspecific databases often results in many unidentified or falsely identified spectra because an overly large or incomplete search space negatively impacts the peptide identification rates[234]. Therefore, for highly-studied systems such as the human gut microbiome, multiple gene catalogs are available, for example, the Integrated Gene Catalog of the human gut microbiome (IGC)[5] or the Unified Human Gastrointestinal Genome (UHGG)[6]. Such catalogs are comprehensive, publicly available, and can be used as a database for gut (fecal) metaproteomics but with the potential risk of having an inaccurate FDR estimation due to their increased size.

Other options are metagenomic or metatranscriptomic databases that are produced by sequencing and assembly of DNA and/or RNA extracted from the same sample. Such meta-omic databases have the advantage to be sample-specific and thus be more fitted for identification but require additional instrumentation and have their own challenges. The most severe challenges are errors occurring during sequencing, assembly, ORF prediction, or binning. Each step in the generation of these sample-specific databases is prone to small errors, but these can accumulate during the full processing path[7].

#### 1.3. Protein grouping methods

A LC-MS/MS experiment will result in peptide spectrum matches (PSMs) obtained from matching experimental spectra with theoretical spectra from theoretical peptides that are extracted from a protein sequence database. Peptides matched in this way can be traced back to their protein of origin, but this additional layer adds a new challenge to data analysis: an identified peptide may be found in multiple proteins. This ambiguity is particularly challenging in metaproteomics, since multiple protein identifications are often plausible.

To address protein identification ambiguity, protein grouping is performed. A protein group aims to combine proteins based on shared peptide identifications. Instead of individual proteins, the group is reported to the user with updated annotation meaningfully derived from the underlying proteins of the group. However, further challenges are encountered during this process: how exactly are proteins grouped and how are taxonomic and functional annotations derived.

To allow protein group comparison, we created groups using the combined peptide evidence of all compared samples. We tested four different protein grouping approaches based on either MPA or PAPPSO and based on protein groups or subgroups

This is reflected in the functional and taxonomic annotation that is derived from these groupings. In the case of proteins of a protein group with conflicting annotation, the annotation for this protein group can become 'various' to reflect the ambiguity. The percentage of groups that are clearly categorized was much higher at the subgroup level than the group level for both PAPPSO and MPA. Since the 'share at least one peptide'-rule for groups is less stringent than the 'share a common peptide set'-rule of subgroups, groups contain more proteins than subgroups, which increases the chance for conflicting assignments leading to a lower percentage of consistently annotated protein groups.

In Suppl. Note 1.3, Figure 1, four different protein grouping methods are illustrated. The PAPPSO grouping (http://pappso.inrae.fr/en/bioinfo/xtandempipeline/) uses two levels, the group and the subgroup, and applies the rule of parsimony to remove proteins without distinct peptide evidence after the grouping algorithm is performed. The MetaProteomeAnalyzer (MPA) uses two major grouping methods that are based on peptides: the “shared peptide” rule and the “shared peptide set” rule, which correspond to the PAPPSO group and subgroup respectively. MPA does not apply the rule of parsimony; instead all available proteins are used.

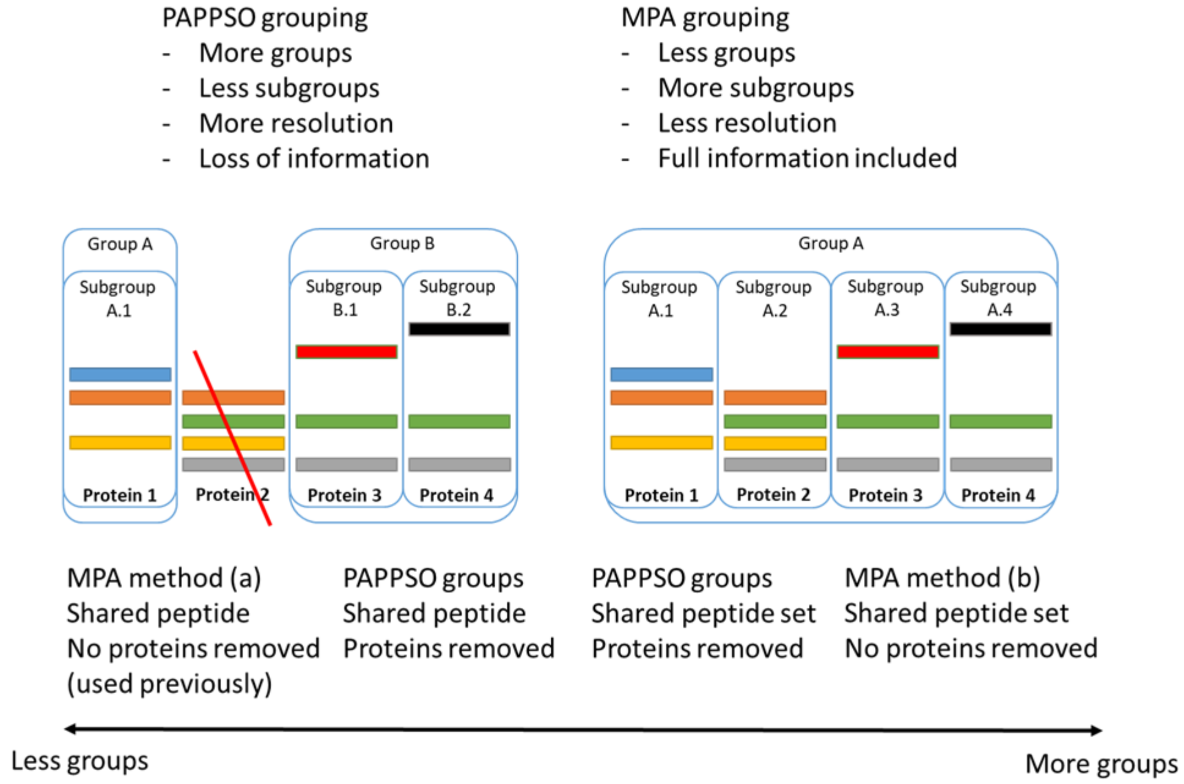


**Suppl. Note 1.3, Figure 1. Protein grouping methods.** Illustration of PAPPSO and MPA protein grouping methods and their differences. For the sake of easy comparison MPA rule “shared peptide” is called “group” and MPA rule “shared peptide set” is called “subgroup”, analogous to PAPPSO. For PAPPSO grouping, a protein is removed after grouping, if it is not identified by at least one peptide that is not found in other identified proteins. However, proteins with the exact same set of peptides are retained. The figure also shows the expected trends for the number and of groups.

To analyze differences in protein grouping strategies, the PSMs of all SIHUMIx samples and all GUT samples were combined. For both, all proteins that contain a given PSM were extracted from the appropriate FASTA file using a Java script (github.com/metaproteomics/CAMPI). From this data, protein groups were created with PAPPSO and MPA. For PAPPSO grouping, the appropriate xml format was used as the exchange format[8]. For MPA grouping the grouping code was directly integrated into the Java scripts. The PSM lists still contained peptides with length six, which proved to be detrimental to protein grouping, by creating large groups of unrelated proteins. Therefore a second analysis was performed excluding peptides of length 6.

The total number of created groups and subgroups and their underlying protein/peptide and psm count are shown in **Supplementary Table 4**.

#### 1.4. Unipept analyses

In the advanced Unipept analysis, which was used with SIHUMIx since the sample composition was known, we used the following steps (scripts available on GitHub):

- For each full peptide, including missed cleavages:
    - Replace J with L (Unipept can't handle "J" as input)
    - run pept2taxa_advanced, returns intersection_peptides.csv
    - run pept2lca_advanced (equate I/L), returns intersection_lca.csv
    - run unipept_advanced_analysis_SIHUMIx.py to establish the level of lca

Example situation 1: if a certain peptide is found in *Clostridium butyricum* and two species unrelated to SIHUMIx; then we can safely remove the other two and calculate the SIHUMIx-specific LCA based on the only SIHUMIx species left, i.e. reassign the LCA to *Clostridium butyricum*.

Example situation 2: if a peptide is assigned to *Clostridium butyricum*, *Bacteroides thetaiotaomicron* and an unrelated SIHUMIx species, the LCA calculation is based on *Clostridium butyricum* and *Escherichia coli*; so the true LCA can be found, but it will not be assigned on species level.

### 2. Supplementary Figures

#### 2.1 Identification: Peptides and Proteins groups comparison

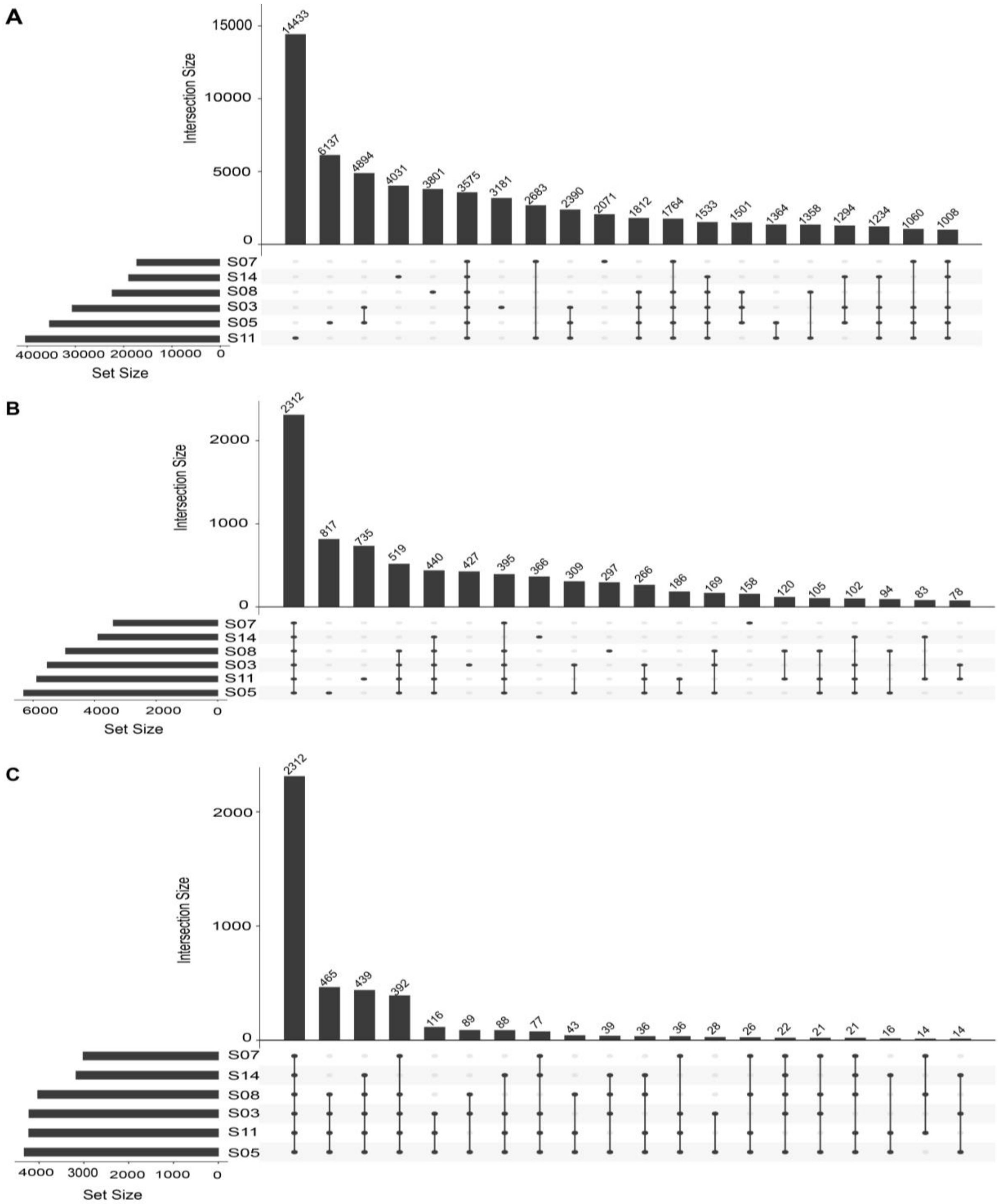


**Suppl. Figure 1. Complete UpSet plot comparison for the SIHUMIx dataset.** The different panels show the comparison at the levels of A) peptides, B) protein subgroups and C) top 50% of the protein subgroups between sample preparations. Unlike Figure 4, the UpSet plot shows the 20 most abundant intersections, regardless of the intersection composition. For Peptides, the unique intersections (size=1) are the most prominent, while they entirely disappear for the top 50% protein subgroups. S03 and S05 are almost identical laboratory workflow, differing in LC gradient length, and the intersection between S03 and S05 is large for the peptides.

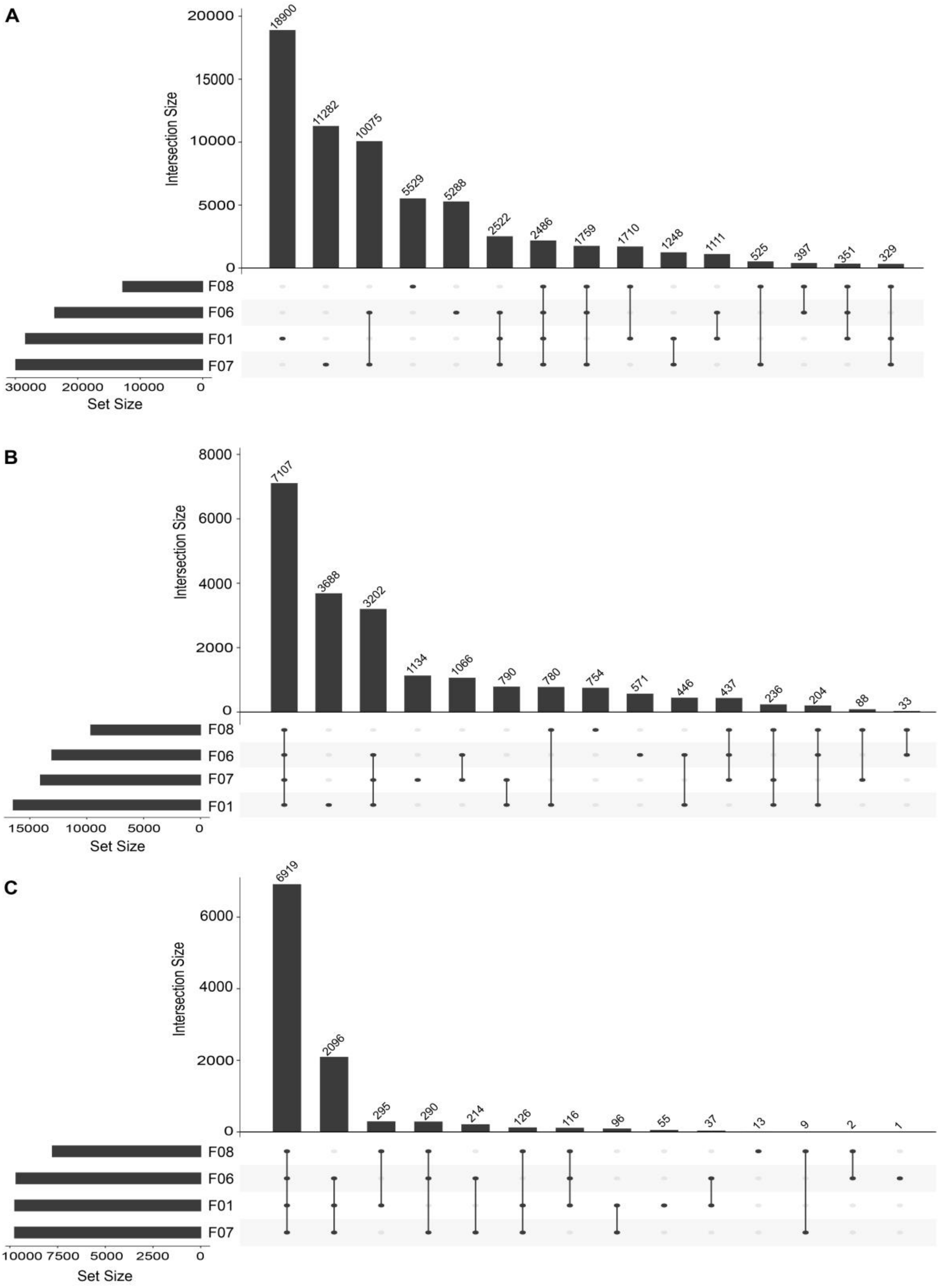


**Suppl. Figure 2. Complete UpSet plot comparison for the FECES dataset.** The different panels show the comparison at the levels of A) peptides, B) protein subgroups and C) top 50% of the protein subgroups between sample preparations. Unlike Figure 5, the UpSet plot shows the 20 most abundant intersections, regardless of the intersection composition. For Peptides, the unique intersections (size=1) are the most prominent, while they entirely disappear for the top 50% protein subgroups. F06 and F07 are almost identical laboratory workflow, one of them utilizing fractionation, and the intersection between F06 and F07 is large for the peptides.

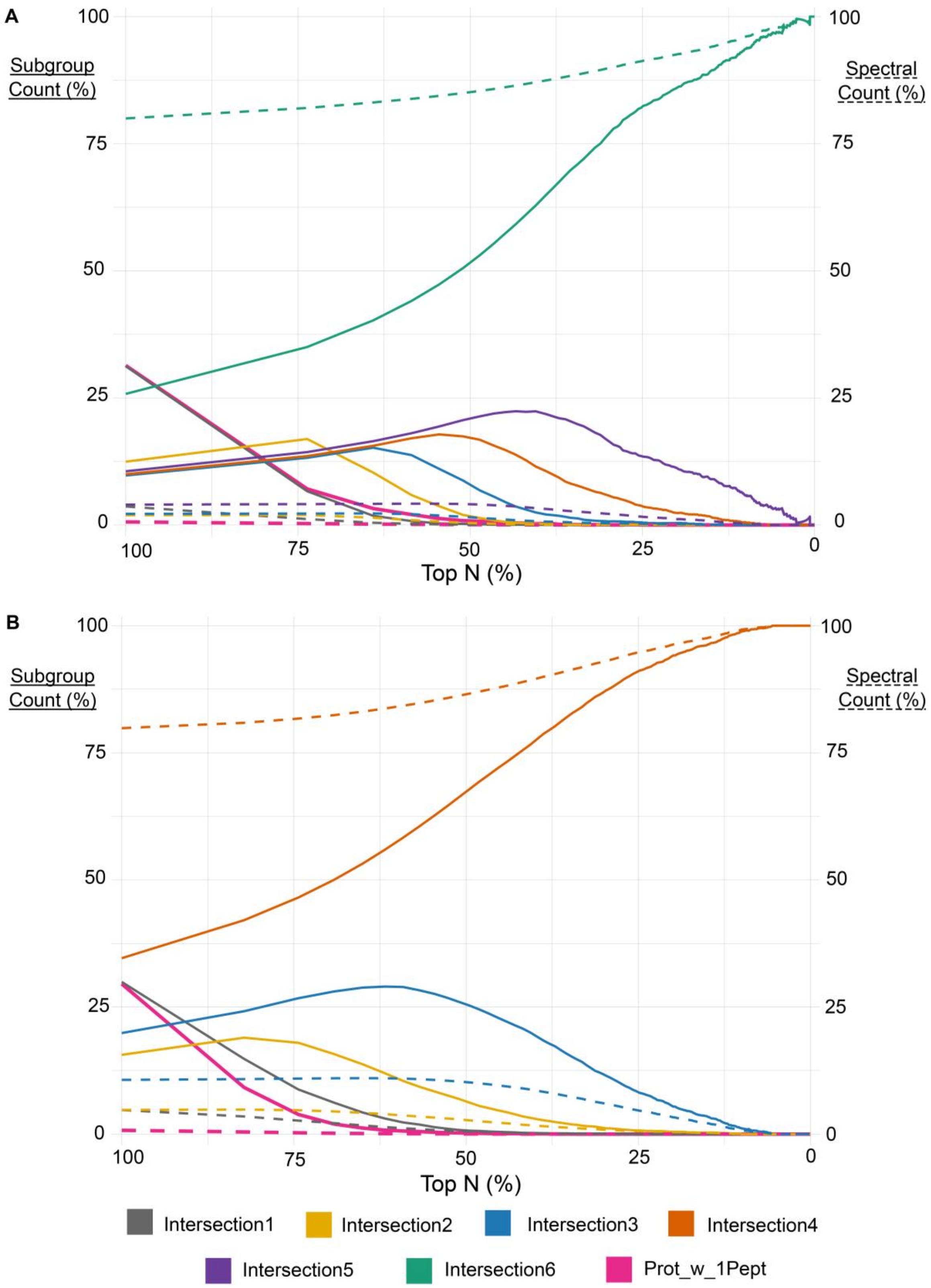


**Suppl. Figure 3. Distribution of protein subgroups for different cross-samples overlaps over the top N percent of the datasets.** The Top-N-% protein subgroups (x-Axis) for SIHUMIx (A) and FECES (B) using the group count (solid lines) and the weighted spectral count (dashed lines) for different cross-sample overlaps (intersection 1-6). The Top-N (%) subgroups are the N-% subgroups (x-Axis) based on spectral abundance. Therefore Top-100% means all data is included and Top-50% means the 50% of subgroups with the highest spectral count are selected. The solid lines represent simple subgroup count for the given categories, while the dashed lines represent weighted spectral count for the subgroups summed over the given category. The categories (Intersection 1-6) represent the agreement on the presence of subgroups as intersection sizes also shown in UpSet plots. This means Intersection 1 counts subgroups (or spectra) for all intersections of size 1 (only found in one analysis). Furthermore, the complete sample overlap corresponds to the intersection 4 for the fecal sample and the intersection for SIHUMIx (this means all analyses find these subgroups). Finally, an additional category for protein subgroups with only one peptide is added (Prow_w_1Pept).

#### 2.2 Protein grouping methods

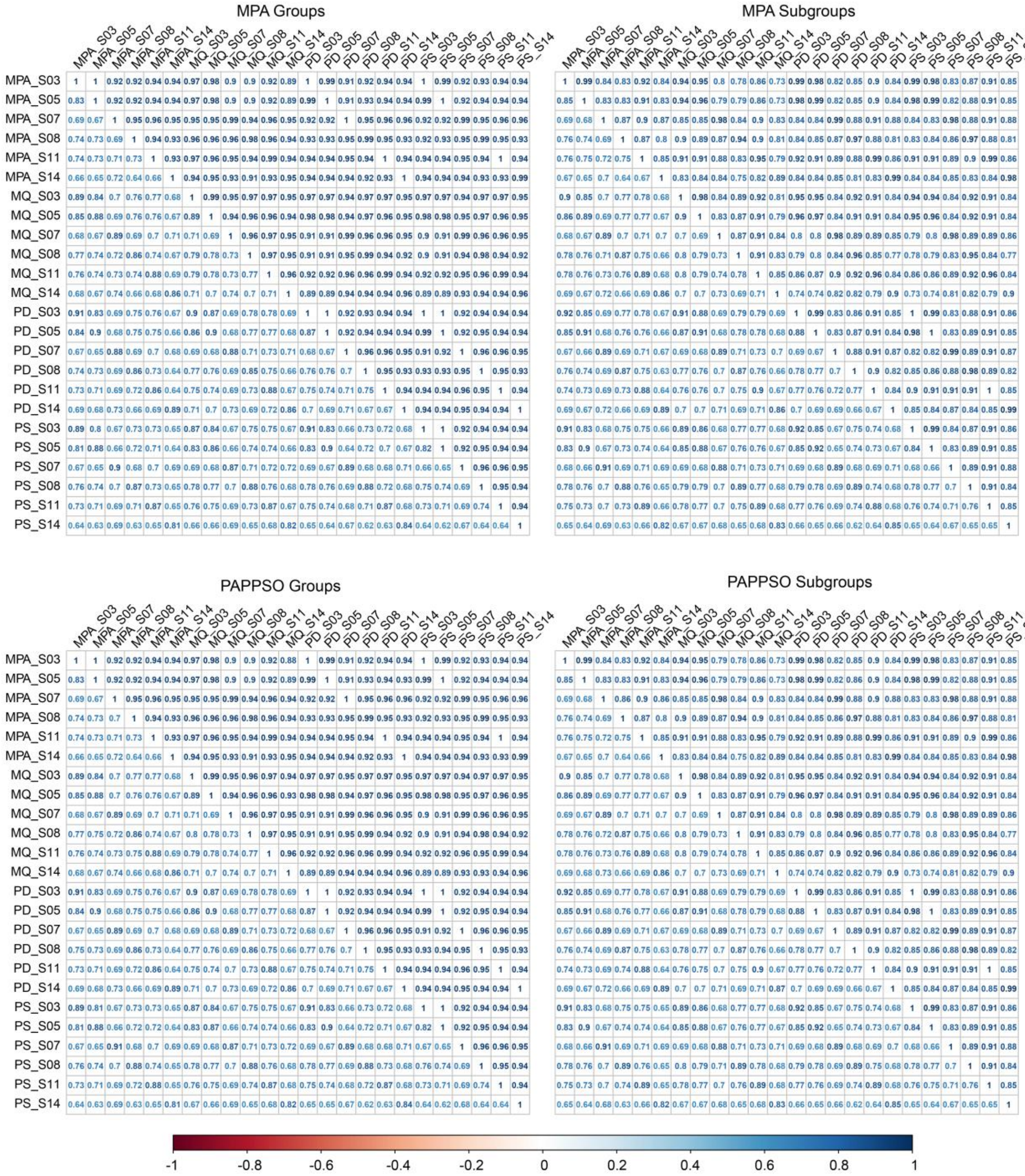


**Suppl. Figure 4. SIHUMIx correlation plots for different grouping methods.** Correlation matrices showing sample-to-sample correlation coefficients for the different grouping methods tried for SIHUMIx. For each plot, the upper and lower triangle show Pearson and Spearman correlation coefficient, respectively. Diagonal lines of higher correlation appear, because they are correlations for the same laboratory workflow for different bioinformatic pipelines, indicating a smaller differences arising from bioinformatic pipelines compared to laboratory workflows.

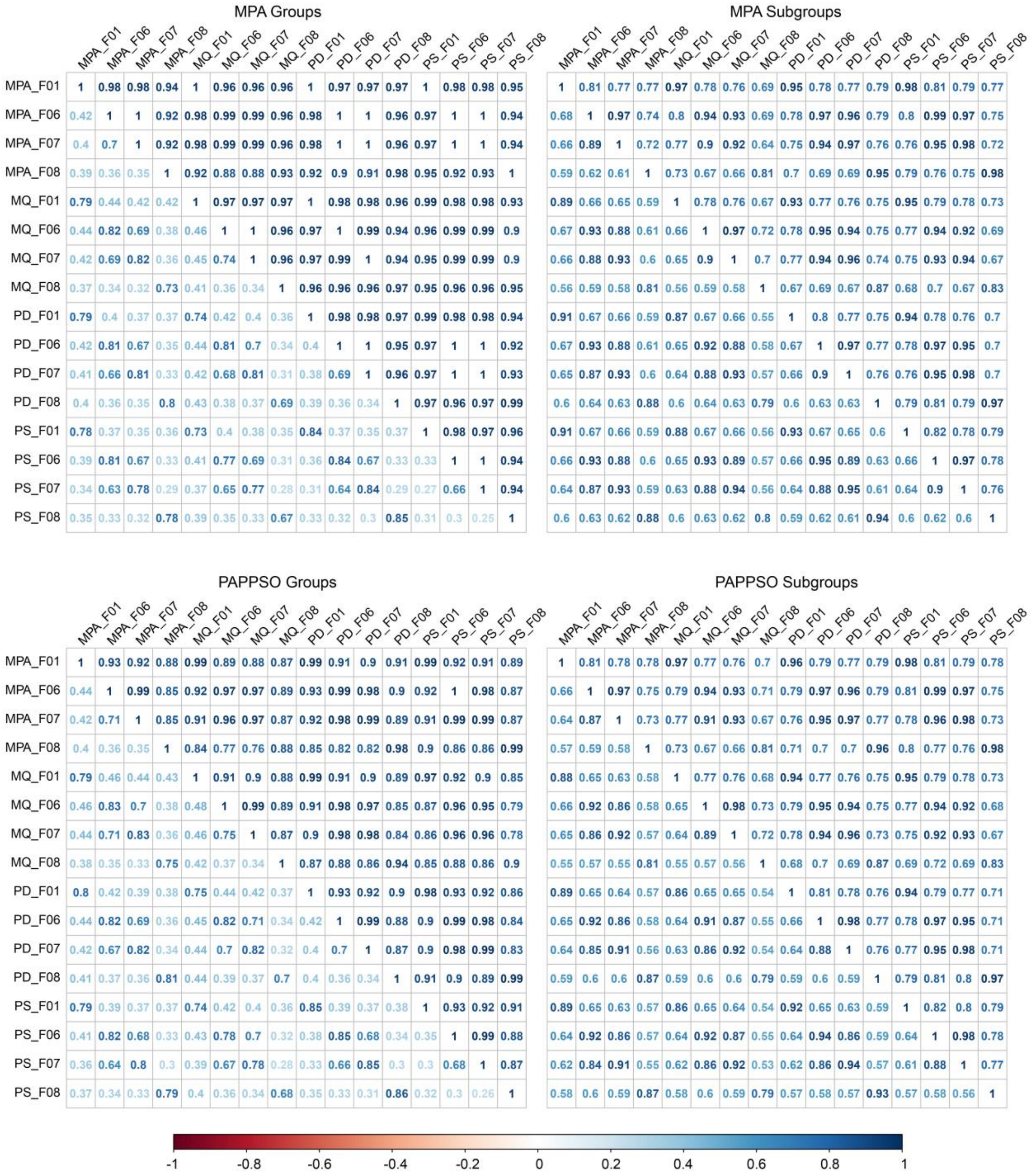


**Suppl. Figure 5. FECES correlation plots for different grouping methods.** Correlation matrices showing sample-to-sample correlation coefficients for the different grouping methods tried for the fecal data set. For each plot, the upper and lower triangle show Pearson and Spearman correlation coefficient, respectively. Diagonal lines of higher correlation appear, because they are correlations for the same laboratory workflow for different bioinformatic pipelines, indicating a smaller difference arising from bioinformatic pipelines compared to laboratory workflows.

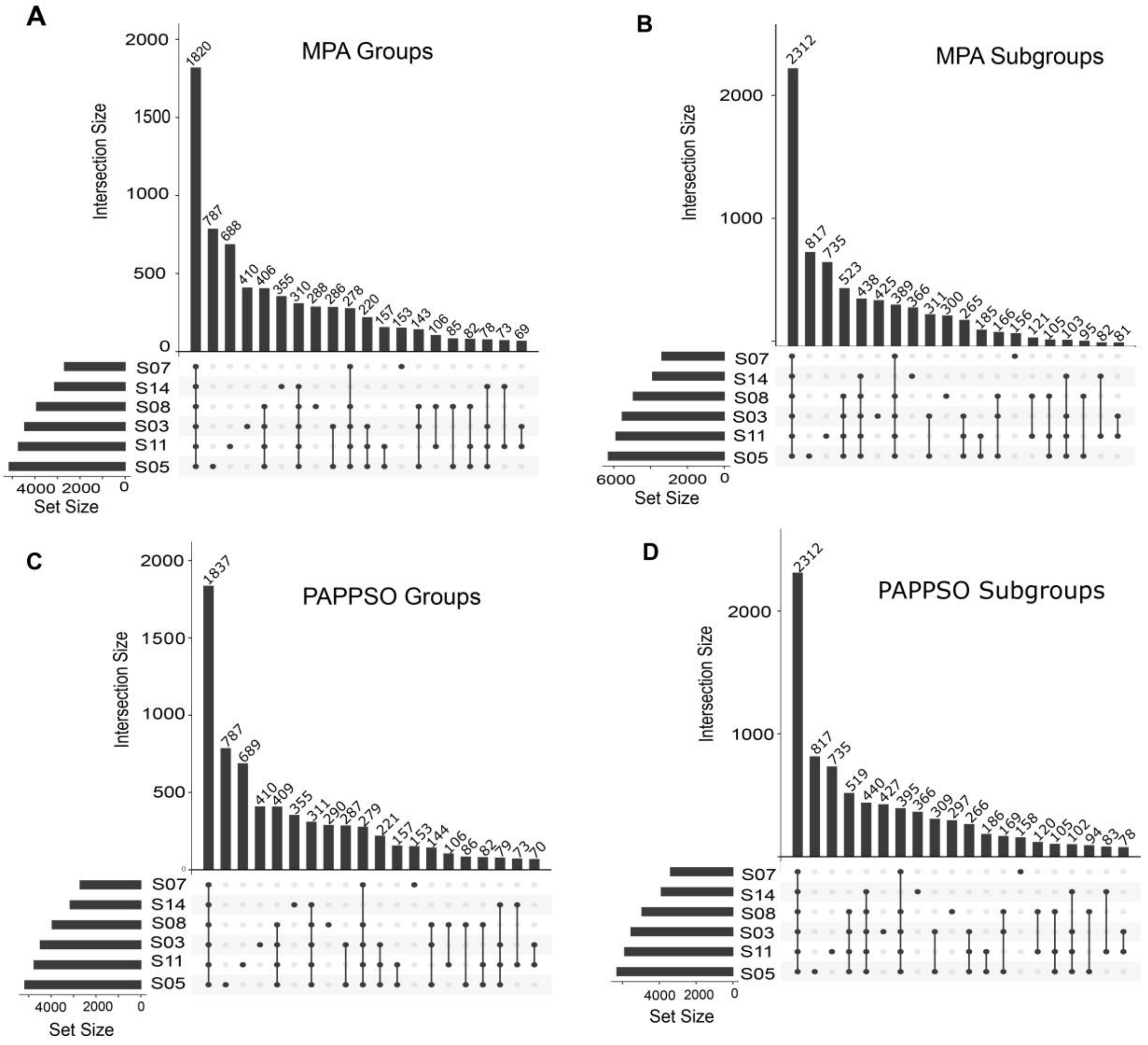


**Suppl. Figure 6. UpSet plot comparison for the SIHUMIx protein grouping.** UpSet plots showing intersection of protein groups (A-C) and subgroups (B-D) for PAPPSO (C-D) and MPA grouping (A-C). All four methods produce highly similar results, and are in line with Suppl. Figure 1B and 2B. Some differences are observable between groups and subgroups, in particular in the amount of subgroups per intersection. Only subtle differences can be observed, between PAPPSO and MPA for both groups and subgroups.

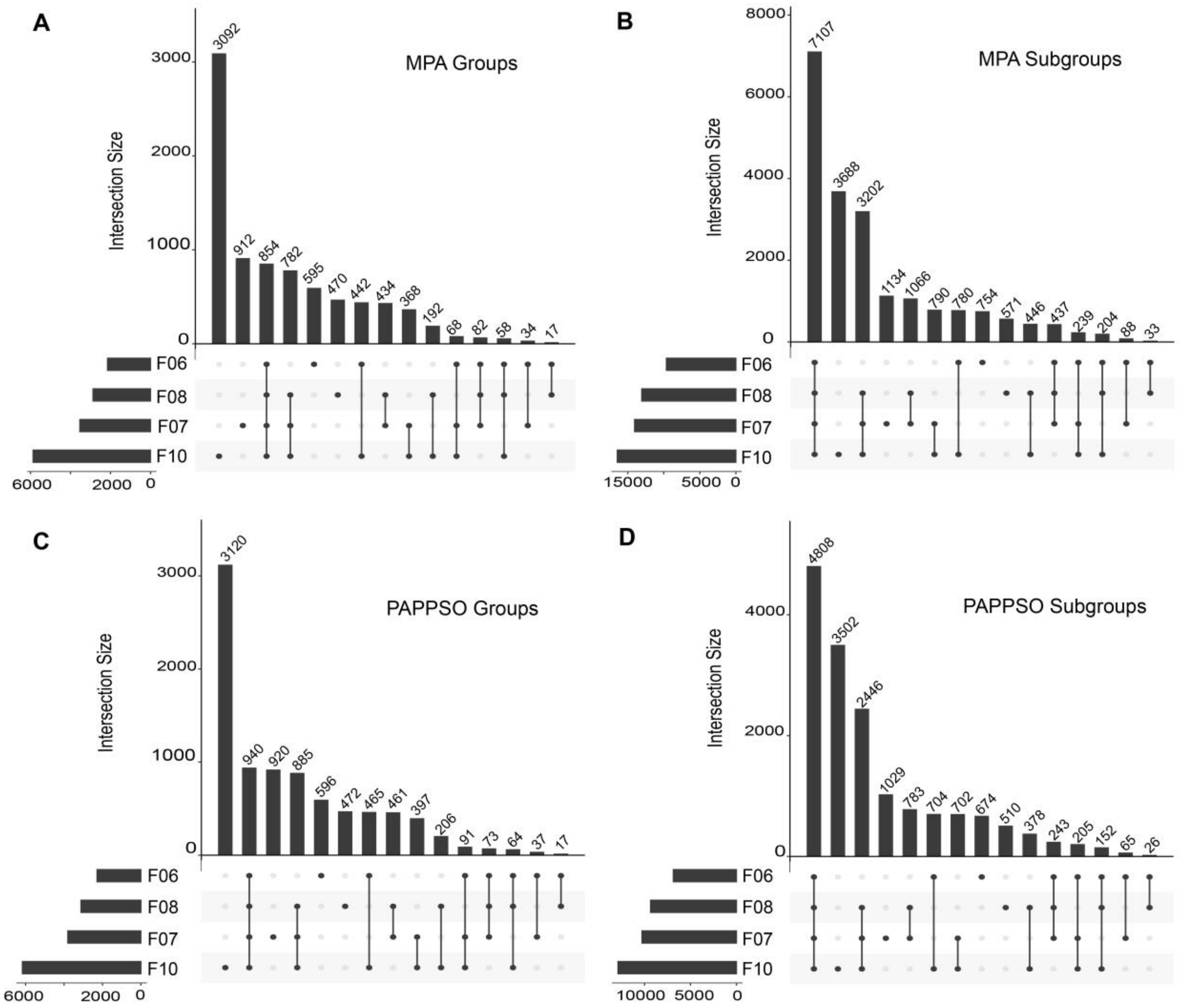


**Suppl. Figure 7. UpSet plot comparison for the FECES protein grouping..** UpSet plots showing intersection of protein groups (A-C) and subgroups (B-D) for PAPPSO (C-D) and MPA grouping (A-C). Groups or subgroups are similar for PAPPSO and MPA in terms of relative size of intersections, but diverge in intersection size. Groups are very different from subgroups, with sample F10 producing a high amount of unique protein groups.

### 2.3 Taxonomic Analyses: Omics Comparison

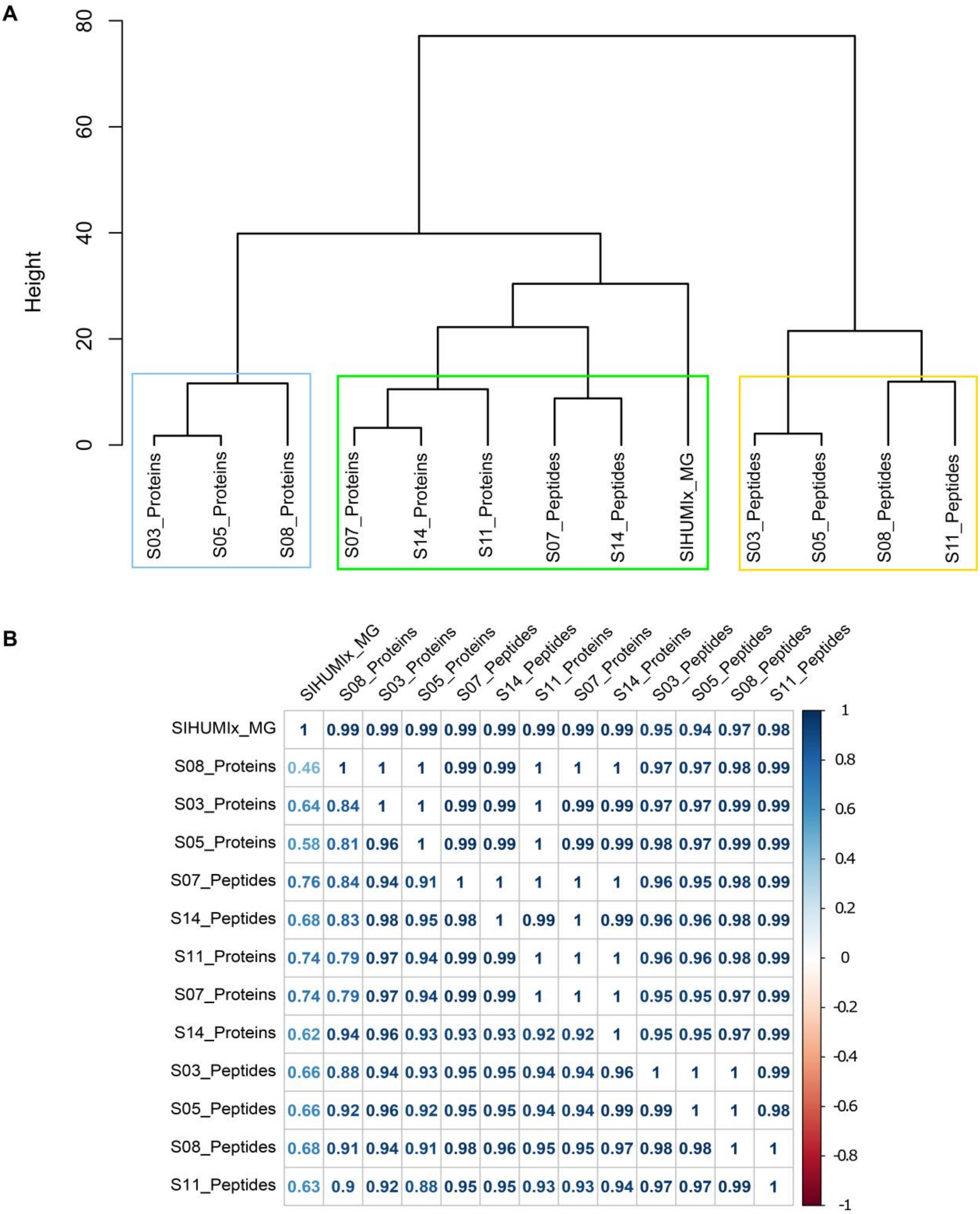


| | SIHUMIx_MG | S08_Proteins | S03_Proteins | S05_Proteins | S07_Peptides | S14_Peptides | S11_Proteins | S07_Proteins | S14_Proteins | S03_Peptides | S05_Peptides | S08_Peptides | S11_Peptides |
|---|---|---|---|---|---|---|---|---|---|---|---|---|---|
| SIHUMIx_MG | 1 | 0.99 | 0.99 | 0.99 | 0.99 | 0.99 | 0.99 | 0.99 | 0.99 | 0.95 | 0.94 | 0.97 | 0.98 |
| S08_Proteins | 0.46 | 1 | 1 | 1 | 0.99 | 0.99 | 1 | 1 | 1 | 0.97 | 0.97 | 0.98 | 0.99 |
| S03_Proteins | 0.64 | 0.84 | 1 | 1 | 0.99 | 0.99 | 1 | 0.99 | 0.99 | 0.97 | 0.97 | 0.99 | 0.99 |
| S05_Proteins | 0.58 | 0.81 | 0.96 | 1 | 0.99 | 0.99 | 1 | 0.99 | 0.99 | 0.98 | 0.97 | 0.99 | 0.99 |
| S07_Peptides | 0.76 | 0.84 | 0.94 | 0.91 | 1 | 1 | 1 | 1 | 1 | 0.96 | 0.95 | 0.98 | 0.99 |
| S14_Peptides | 0.68 | 0.83 | 0.98 | 0.95 | 0.98 | 1 | 0.99 | 1 | 0.99 | 0.96 | 0.96 | 0.98 | 0.99 |
| S11_Proteins | 0.74 | 0.79 | 0.97 | 0.94 | 0.99 | 0.99 | 1 | 1 | 1 | 0.96 | 0.96 | 0.98 | 0.99 |
| S07_Proteins | 0.74 | 0.79 | 0.97 | 0.94 | 0.99 | 0.99 | 1 | 1 | 1 | 0.95 | 0.95 | 0.97 | 0.99 |
| S14_Proteins | 0.62 | 0.94 | 0.96 | 0.93 | 0.93 | 0.93 | 0.92 | 0.92 | 1 | 0.95 | 0.95 | 0.97 | 0.99 |
| S03_Peptides | 0.66 | 0.88 | 0.94 | 0.93 | 0.95 | 0.95 | 0.94 | 0.94 | 0.96 | 1 | 1 | 1 | 0.99 |
| S05_Peptides | 0.66 | 0.92 | 0.96 | 0.92 | 0.95 | 0.95 | 0.94 | 0.94 | 0.99 | 0.99 | 1 | 1 | 0.98 |
| S08_Peptides | 0.68 | 0.91 | 0.94 | 0.91 | 0.98 | 0.96 | 0.95 | 0.95 | 0.97 | 0.98 | 0.98 | 1 | 1 |
| S11_Peptides | 0.63 | 0.9 | 0.92 | 0.88 | 0.95 | 0.95 | 0.93 | 0.93 | 0.94 | 0.97 | 0.97 | 0.99 | 1 |

**Suppl. Figure 8.** Dendrogram based on Manhattan distance for SIHUMIx sample and corresponding correlations values. The upper triangle is based on Pearson correlation and the lower triangle on Spearman’s. Empty cases indicate non-significant correlations.

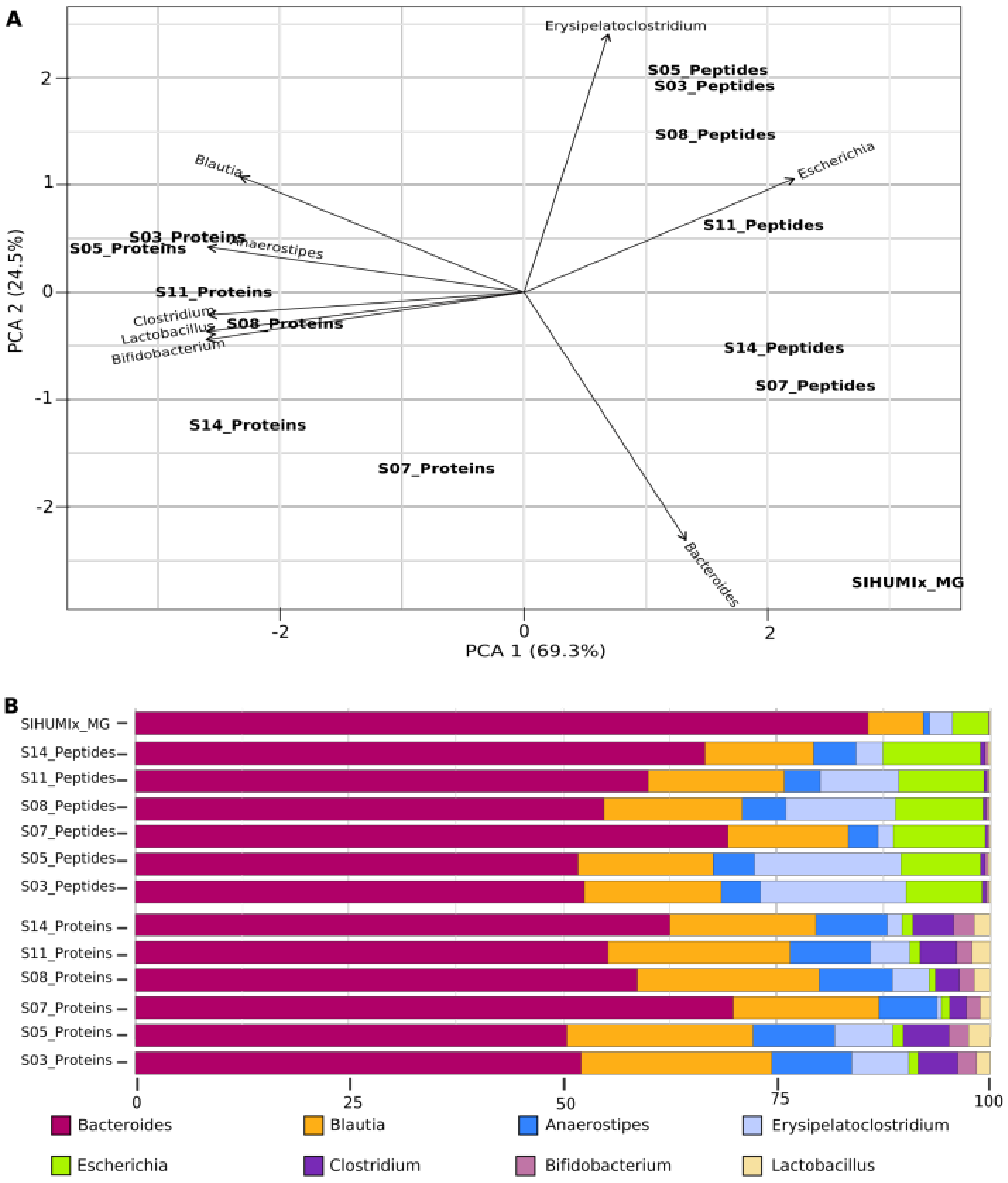


**Supplementary Figure 9. Comparisons of community composition for SIHUMIx at the genus level.** The upper panel shows PCA clustering of the results (A). Different approaches and tools used for taxonomic annotation (mOTU2, Unipept and Prophane) are indicated in the label. Clusters (k=3) were calculated using manhattan distance and are represented by blue, yellow, and green. Features not annotated at species level were considered unclassified and discarded for PCA calculation. Unclassified features accounted for 24.2% and 69.9% of data for peptide and protein subgroup levels. Variables driving differences between samples are represented by black arrows. The lower panel details taxonomic profiles of each sample as bar plots (B).

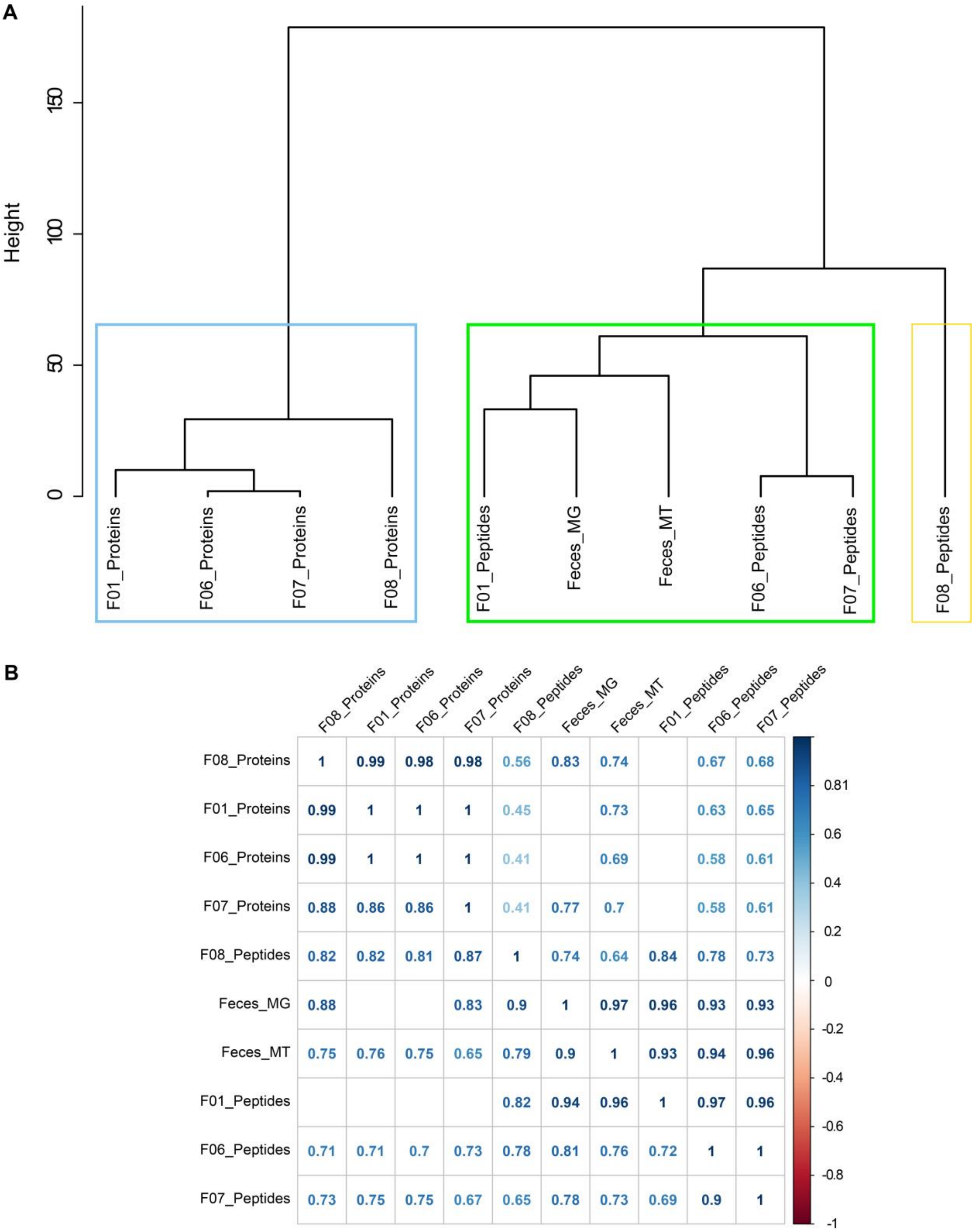


**Suppl. Figure 10.** Dendrogram based on Manhattan distance for fecal sample and corresponding correlations values. The upper triangle is based on Pearson correlation and the lower triangle on Spearman's. Empty cases indicate non-significant correlations.

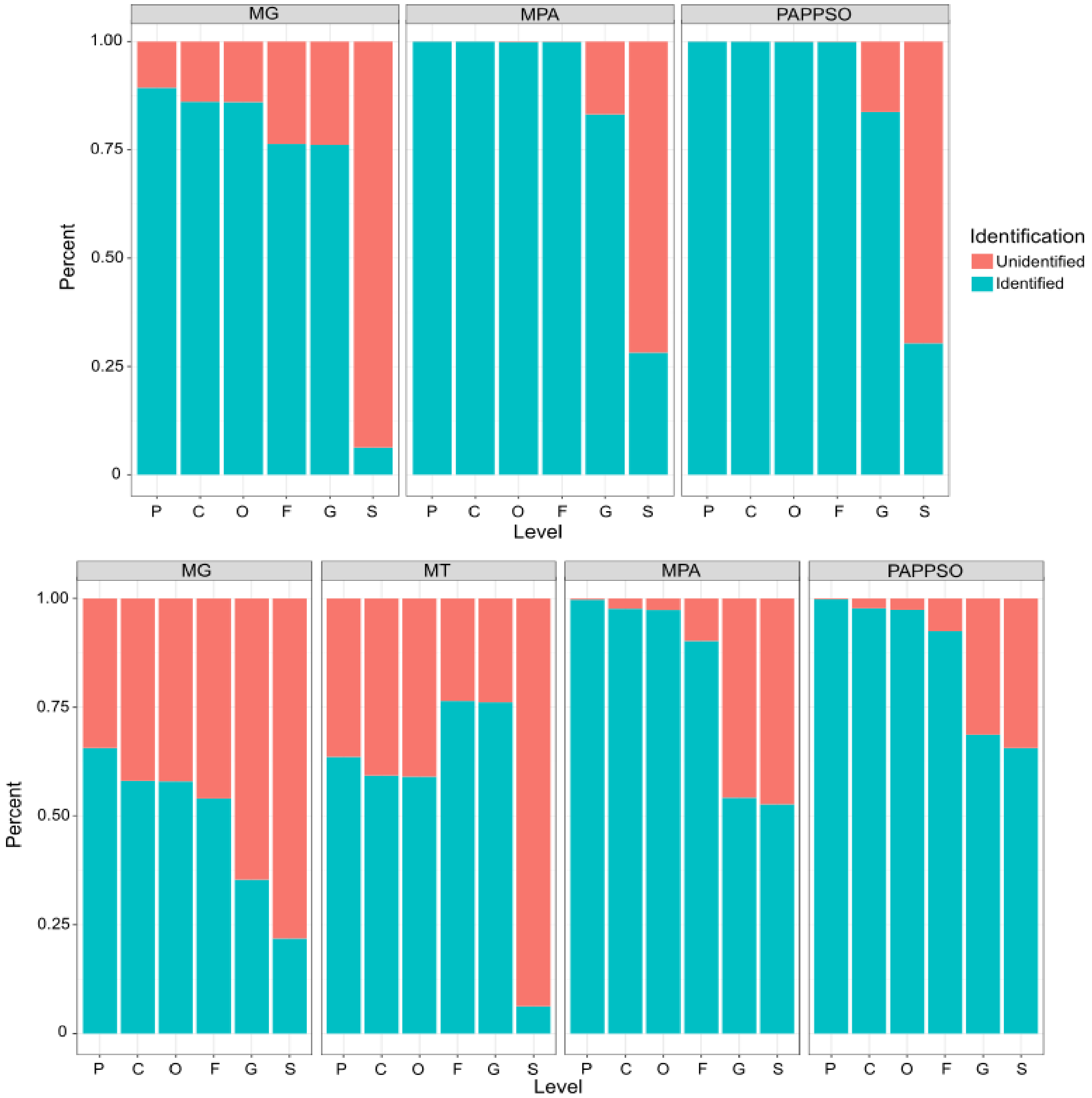


**Suppl. Figure 11 A/B: Taxonomic resolution across omics domains for SIHUMIx (a) and the Fecal (b) samples.** The stacked barplots represent the percentage of metagenomic (MG) and metatranscriptomic (MT) reads or identified proteins (MPA and PAPPSO) that can be annotated at the Phylum (P), Class (C), Order (O), Family (F), Genus (G) and Species (S) levels. For metaproteomics, percentage of annotation starts to decrease at the Genus level and becomes very low at the Species level. For the complex fecal sample (**b**), PAPPSO grouping allows better taxonomic resolution as already described in the supplementary note 1.3. For metagenomics and metatranscriptomics, the lower resolution is also higher at the species level but already appears at the Phylum level. This is mostly due to the method used to allow such comparison. Indeed instead of using the marker gene-based method, we annotated every single read with Kraken2, while the metaproteomics approach only focused on identified proteins. This important difference in size of the data and the fact that metaproteomics analyses a subset of it can justify the differences observed here.

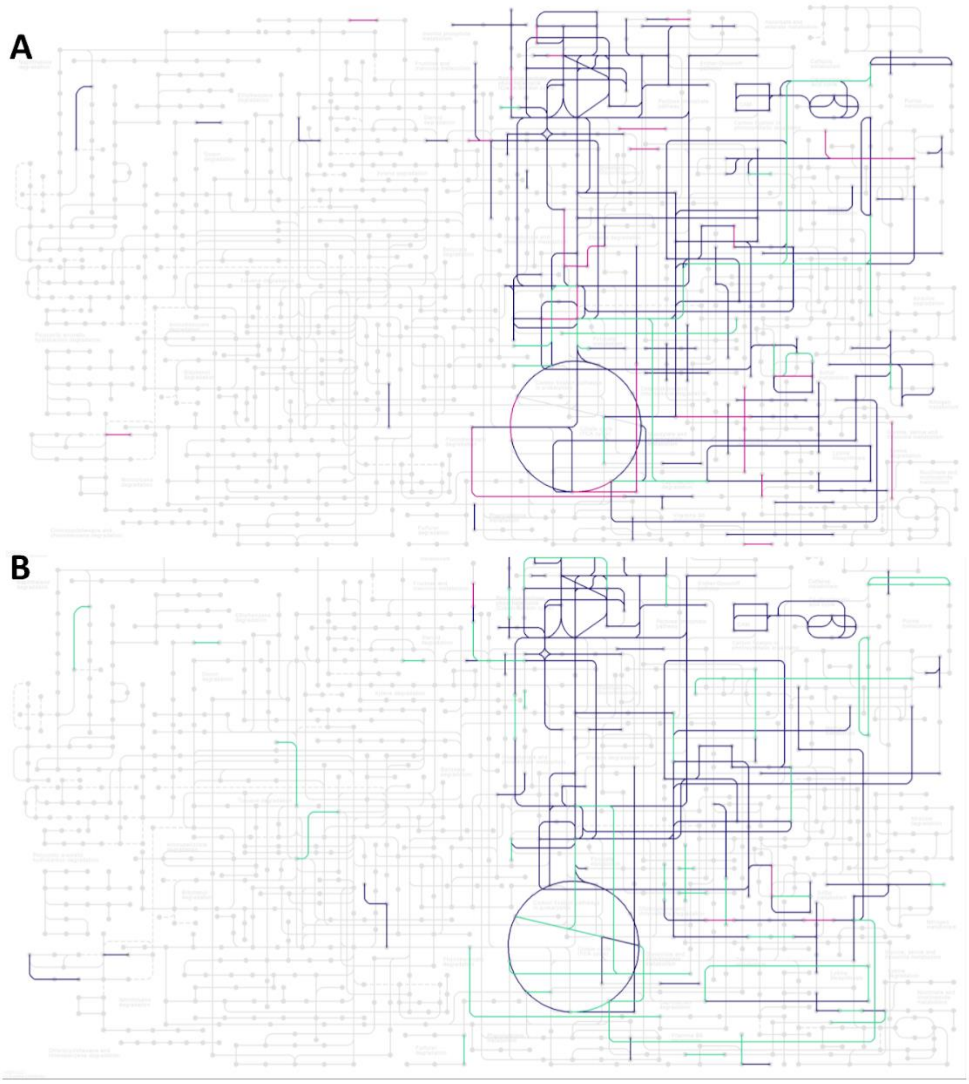


**Suppl. Figure 12. Mapping of the KEGG ID onto the Microbial metabolism in diverse environments KEGG map for SIHUMIx (A) and FECAL (B) samples.**

The most (S11/F01) and the least (S07/F06) sensitive approaches are displayed in green and pink, respectively. The overlap is shown in purple. As already shown in the figure 4, each method brings its own uniqueness, with some approaches being deeper than others. For SIHUMIx, S07 has only 1.7% of unique protein subgroups against 8,1% for S11 (Figure 4C). Similar observations can be made in panel A, where almost no reactions are found uniquely in S07 in contrast to S11.

The fecal samples show similar results (panel B), with the small difference that the least sensitive approach (F06) has more uniquely found functions associated as already discussed in figure 4. Indeed, the more complex the sample is, the more marked the differences are between each method.

It is interesting to note that no new pathways were identified by the most sensitive methods in comparison to the least sensitive ones. Instead, similar pathways are identified but with a bigger coverage of reactions. The only exception is for SIHUMIx, where S11 identifies a partial "purine metabolism" that is missed by S07.

Finally, it should be noted that even though there are differences in pathway completeness in terms of identification, those are minimal when the quantification is taken into account. As shown in supplementary figure 3, the uniquely identified protein subgroups for all datasets are usually identified with only one peptide and account for less than 4% of the total spectra.

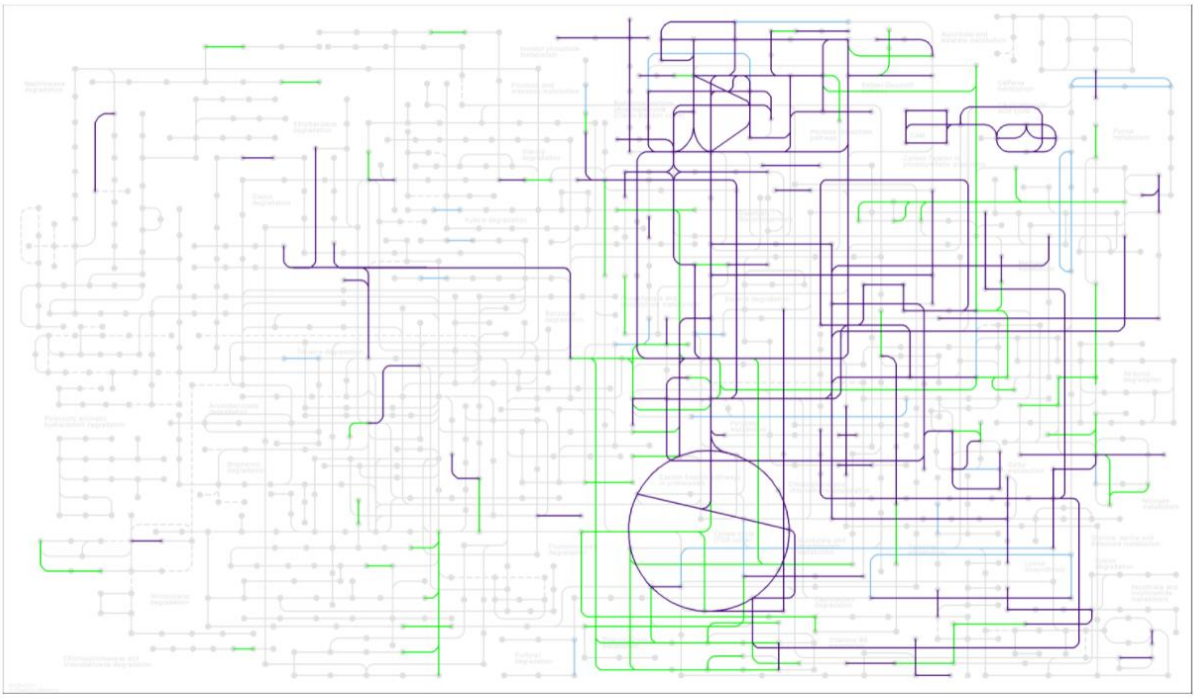

**Suppl. Figure 13. Mapping of the KEGG ID onto the Microbial metabolism in diverse environments KEGG map for SIHUMIx.** Metagenomics, metaproteomics and the overlap are displayed in green, blue and purple, respectively.

As expected, we can observe that the expressed functions found in the metaproteomics data correspond to a subset of those identified in the metagenomics data representing the complete potential of the community and mostly overlap it. Few functions are found only by metaproteomics, but this is due to the different databases used for the SIHUMIx metaproteomic analysis. Indeed, the reference database was used for the search, showing that few functions were missed during the generation of the metagenomic database.

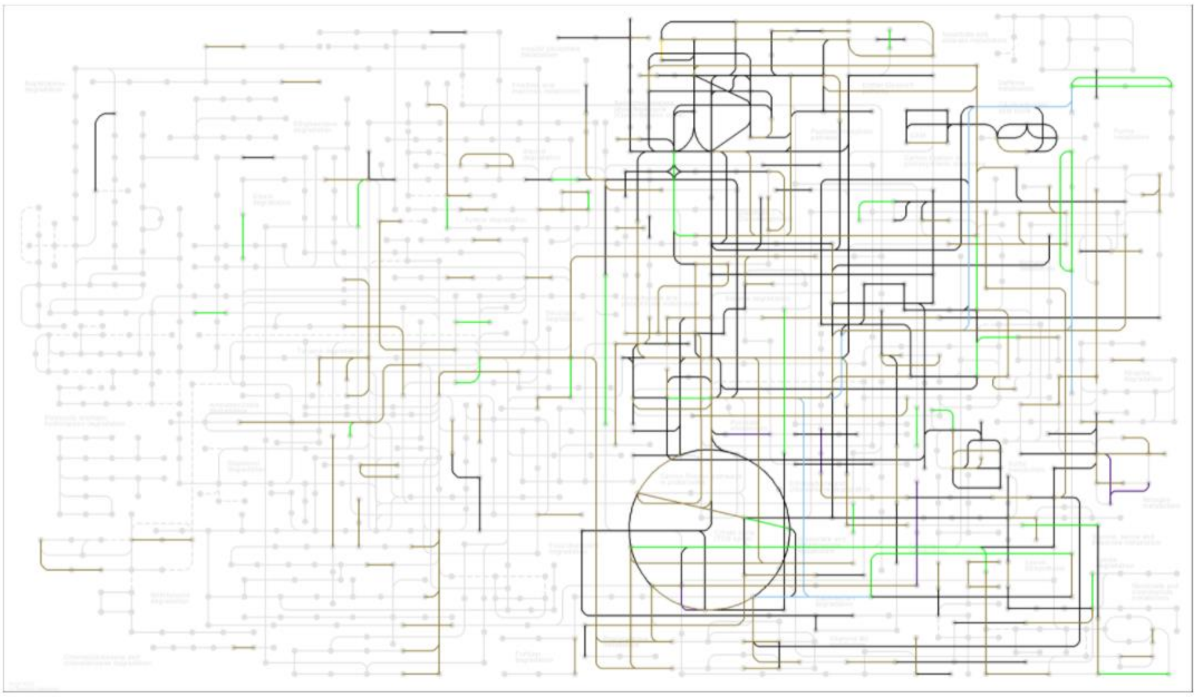

**Suppl. Figure 14. Mapping of the KEGG ID onto the Microbial metabolism in diverse environments KEGG map for Gut.** Metagenomics, metatranscriptomics and metaproteomics are displayed in green, yellow and blue, respectively. Overlap between metagenomics and metatranscriptomics is coloured in brown, overlap between metagenomic and metaproteomics is coloured in purple and overlap between all omics domains is coloured in black.

Similarly to SIHUMIx, the pathways identified in the metaproteomics dara are mostly a subset of the metagenomic pathways. Interestingly, very few functions were uniquely found via metatranscriptomics (yellow) but a good part of the functions found by metagenomics are covered by metatranscriptomics (brown) but not by metaproteomics. This indicates that the functional potential identified by metatranscriptomics is closer to the one identified by metagenomics than the one from metaproteomics and that the latter is lacking depth of analysis.

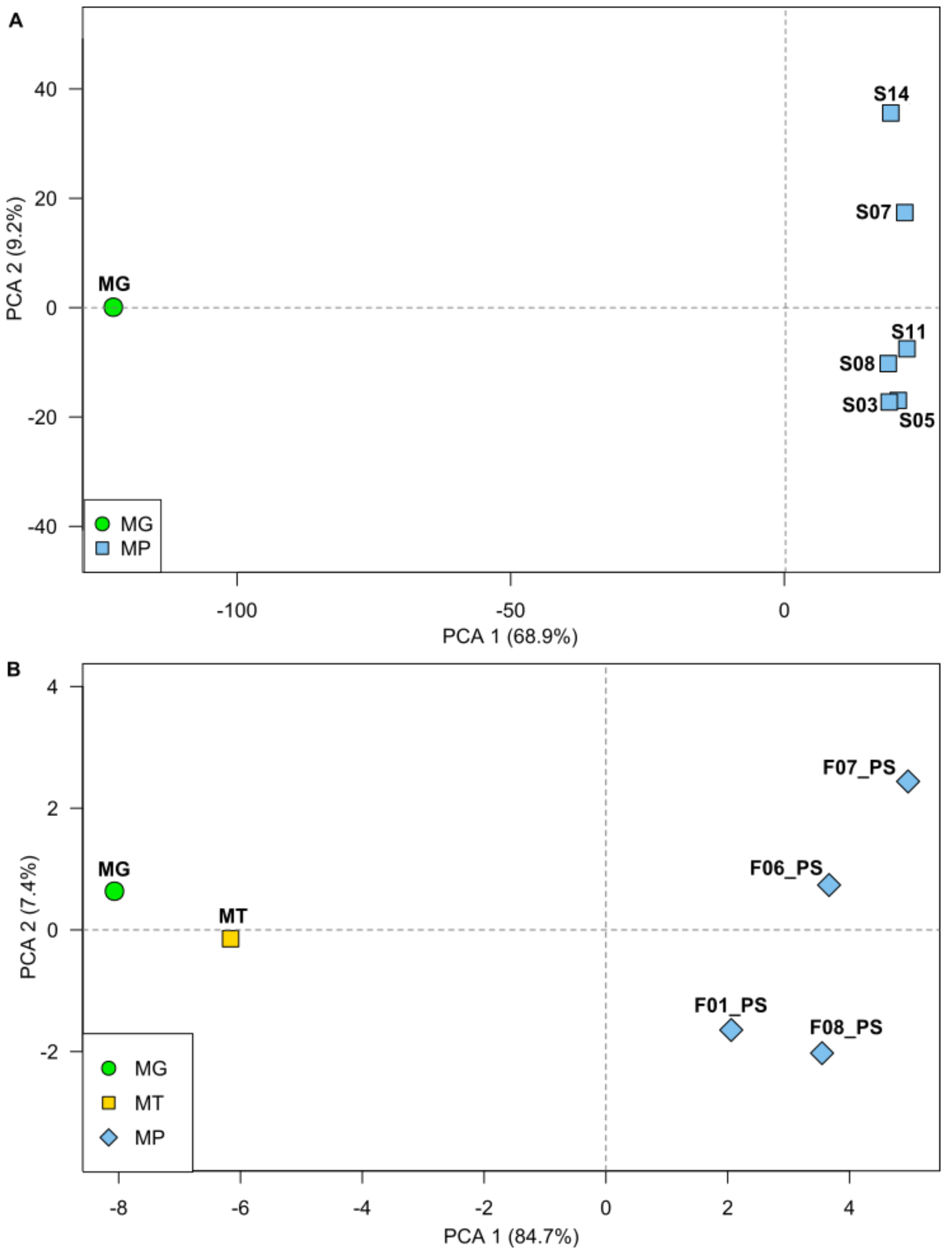


**Suppl. Figure 154. Comparisons of community's functional composition for the SIHUMIx (A) and fecal (B) sample .** The PCA analysis shows a strong separation between metagenomics (green) and the different metaproteomic samples (blue). Metatranscriptomics (yellow) is in between the two other omics domains but very similar to metagenomics.

Here, to go further into the comparison and add the quantitative aspect, we retrieved the read mapping counts for each ORF at the metagenomic/metatranscriptomic level and spectral counting for each protein subgroup at the metaproteomic level. We retrieved the KEGG annotations and compared the omics domains using PCA (Supplementary Figures 14 A and B). Similarly to the PCA plots for the taxonomic profiles, we can observe differences between omics domains, with metagenomics being relatively distant to the other omics domains and metaproteomics and metatranscriptomics being relatively close to each other. As expected, the differences at the functional level are much stronger than at the taxonomic level. Indeed, when a taxon switches metabolism, no differences will be observed in the taxonomic profile of the community, but the functional profile will be affected, thus leading to bigger differences at the functional level.
